# Curcumin attenuates microglia-mediated chronic neuropathic pain through CDK5 /p35 signaling pathway

**DOI:** 10.1101/2024.11.13.623498

**Authors:** Fei Tong, li Zhao, Yue Yang, ZhuoYue Zhang, Lu Liu, YaXuan Wu, XuanYi Di, ZiWen Zhang, XiaoXia Xu, YuanLi Zhang, Yue Shen, liang Yu, lei Zhang, YongXing Yao, HongHai Zhang

## Abstract

Curcumin is a phenolic compound derived from turmeric, one of the main ingredients of curry powder, which is widely used for its antioxidant, anti-inflammatory and immunomodulatory effects. Curcumin has been reported to help relieve pain, such as neuropathic pain caused by injury or disease, but the specific mechanism of its antinociceptive effect on pathological pain is unclear. Cyclin-dependent kinase 5 (Cdk5) is a key control point for the release of neurotransmitters from presynaptic vesicles. Cdk5 and its activator p35 couple to regulate key signaling cascades, thereby participating in the pain process. In this study, we established a NP model by chronic constriction injury (CCI) of the bilateral sciatic nerve in rats and evaluated behavioral hyperalgesia using mechanical and hot and cold tests. Protein expression and distribution were evaluated using western blotting and immunofluorescence. The results showed that Iba-1 and Cdk5/p35 were co-localized in the dorsal horn and dorsal root ganglia, respectively. After CCI, the expression of Cdk5 and p35 was upregulated in the dorsal horn and dorsal root ganglia, while intraperitoneal injection of curcumin significantly reversed the activation of Cdk5/p35 protein and alleviated the hyperalgesia in rats. In addition, the injection of curcumin reduced the co-localization expression of Iba-1 and Cdk5/p35, indicating that curcumin inhibited the activation of Cdk5/p35 protein in the dorsal horn and dorsal root ganglia, thereby affecting the activation of microglia, thereby having a destructive effect on the neuronal cell plasticity and synaptic structure remodeling in the development of NP. Our study provides new evidence that Cdk5/p35 in the dorsal horn and dorsal root ganglia is related to the occurrence of NP, introduces microglia as the basis for the long-term maintenance of NP, and provides insights into the molecular mechanisms involved in the analgesic effect of curcumin.

**Graphical Abstract:** 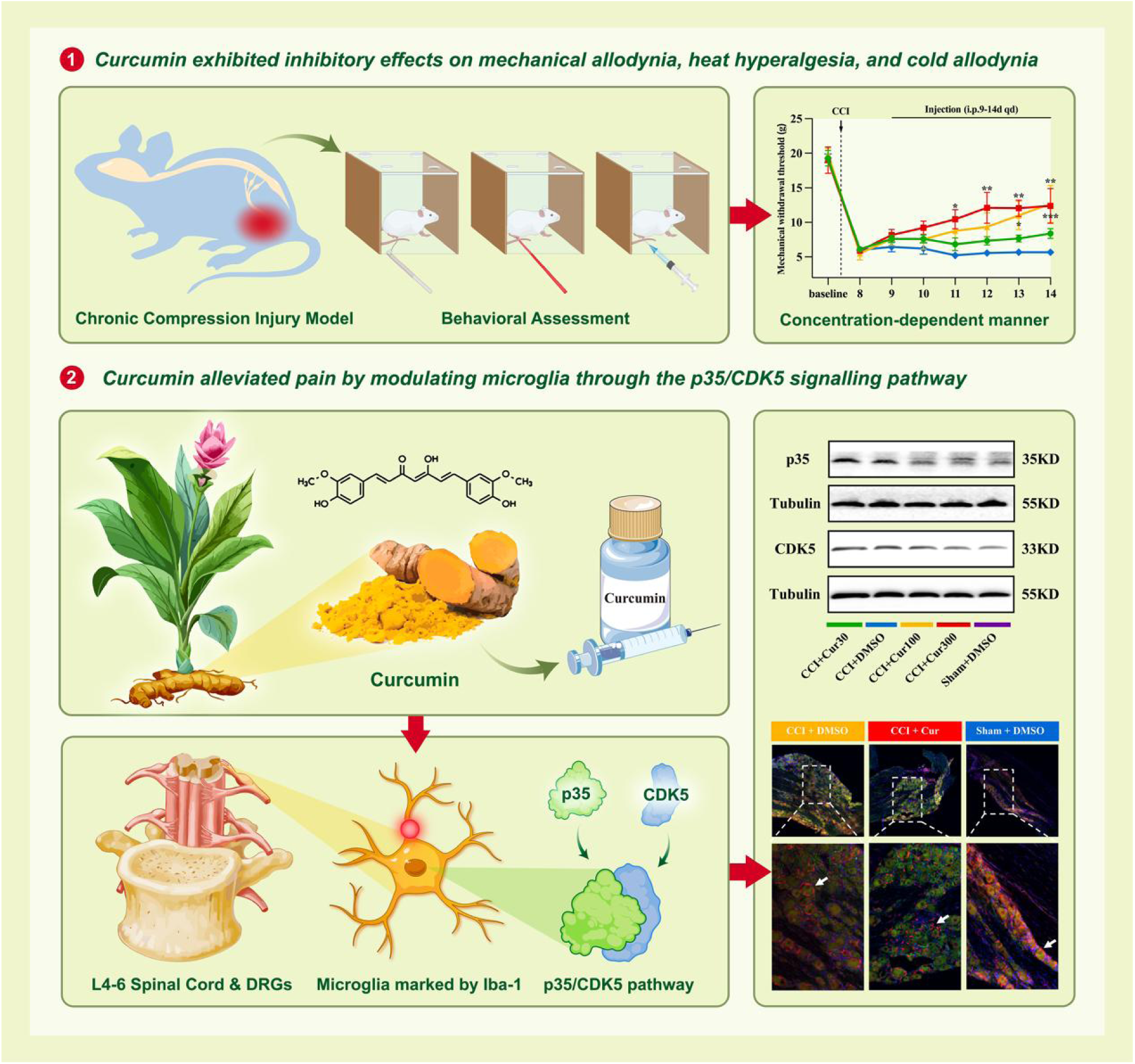

## 1 Introduction

Neuropathic pain (NP), stemming from lesions or injuries of the peripheral or central nervous systems, and it is commonly accompanied by paroxysmal pain, allodynia, and hyperalgesia(An et al., 2021). Accounting for 15%–25% of chronic pain, neuropathic pain significantly affects the quality of life and imposes a heavy burden on public health(Cohen et al., 2021). Although some hypotheses suggest that the occurrence of NP may be related to the changes in neuronal plasticity at various nerve levels, the specific mechanism of NP still needs to be further explored(Bielopolski et al., 2019). Currently, the drugs commonly used in the clinical treatment of neuropathic pain are not effective and have many side effects. Therefore, seeking for drugs with better effect and less side effects for neuropathic pain has become one of the hot topics in the current research on neuropathic pain.

Curcumin, a phenolic compound extracted from Curcuma longa (commonly known as turmeric), serves as a primary constituent in curry powder(Sun et al., 2018). As a naturally occurring phytochemical, it has a variety of pharmacological activities such as anti-inflammatory, anti-tumor, anti-angiogenic anti-angiogenesis, anti-metastasis and anti-multidrug resistance, and is also recognized as a safe substance by GRAS (General Recognition and Safety) of the US FDA (Food and Drug Administration)(Inchingolo et al., 2022). More importantly, curcumin has been reported to have antinociceptive effects in sciatic nerve injury, spinal cord injury, diabetic neuropathy, alcoholic neuropathy, opioid tolerance or opioid-induced hyperalgesia, chemotherapy-induced peripheral nerve inflammation, pathological pain caused by complete Freund’s adjuvant (CFA) injection, or carrageenan injection(Gao et al., 2022; Kong et al., 2023; Sun et al., 2018; Z. Zhao et al., 2017). As previously discussed, the fundamental etiology of neuropathic pain (NP) involves damage to either the peripheral or central nervous system, resulting from lesions, metabolic disorders, or exposure to substances like chemotherapeutic drugs(Cohen and Mao, 2014). This condition is characterized by heightened sensitivity to painful stimuli (hyperalgesia), the perception of pain in response to non-painful stimuli (allodynia), and spontaneous pain sensations. Meanwhile, curcumin’s analgesic effectiveness is attributed to its diverse mechanisms, which encompass mitigating oxidative stress-induced damage, suppressing inflammatory cytokines, modulating glial cell activation, and inhibiting protein kinase activation, among several others. These mechanisms are intricately linked to the underlying causes of neuropathic pain(Baron et al., 2010). Therefore, there is a huge space to explore the aspects of curcumin in the treatment of pathological pain.

In the dorsal nucleus-spinal cord descending pain inhibition pathway, the amount and duration of neurotransmitter release within the synaptic cleft are critical. The release of neurotransmitters relies on the movement of synaptic vesicles, the rupture of fusion with the synaptic plasma membrane, and the process of continuous neurotransmitter transport in vesicle exocytosis and endocytosis. A key factor in this process is the function of various vesicle proteins(Yan et al., 2019). Recent studies have found that Cdk5/P35 plays an important role in pain(Bardin et al., 2003). Cyclin-dependent kinase 5 (Cdk5), a proline-directed serine/threonine kinase, belongs to the Cdks family, which is implicated in transcription regulation, mRNA processing, and neuronal differentiation. While most Cdk5 exists in a monomeric form lacking kinase activity, it becomes enzymatically active only when bound to P35 or P39. Current research indicates that P35 is more characteristic and possesses a stronger ability to bind and activate Cdk5. Cdk5/P35 plays a pivotal role in the nervous system as a major regulator of neuronal function. It is involved in various processes during development, such as neuronal migration, neurite outgrowth(R et al., 2004), axonal transport, exocytosis, drug addiction(Roufayel and Murshid, 2019), learning and memory(S, 1999), and cognitive functions(Chang et al., 2021). Consequently, Cdk5/P35 is considered a crucial modulator in neurological disorders. Previous studies have shown that the tight binding of Cdk5 to p35 can regulate a series of key signaling cascades that are involved in the pathological process of inflammatory pain(V et al., 2022; Yang et al., 2014; Zhang et al., 2021). In a mouse model of inflammatory pain constructed using carrageenan formalin or complete Freund’s adjuvant (CFA), Cdk5 effectively facilitates the transmission of peripheral nociceptive messages to the dorsal horn of the spinal cord by phosphorylating capsaicin receptors within the dorsal root ganglia (DRG)(V et al., 2022; Yang et al., 2014). At the same time, the activity of Cdk5 in the spinal cord increased significantly, highlighting its central position in pain signaling. Notably, Roscovitine, a selective inhibitor of Cdk5, was able to effectively reduce the thermal pain threshold in the inflammatory pain model, but had no significant effect on mechanical pain, providing clues to the specific regulation of pain types by Cdk5(W. Zhao et al., 2017). Based on the above research results and extensive literature review, we try to hypothesize that CDK5/P35 is deeply involved in the regulation of neuropathic pain, and more importantly, curcumin may play an important role in the occurrence and development of NP by affecting the release of CDK5/P35.

Microglia can be activated by a variety of stimulus signals in the early stage of the pain process, and then proliferate and release a large number of inflammatory mediators, regulate synaptic plasticity, and enhance the conduction of pain signals in the spinal cord(Grace et al., 2014; Ji et al., 2013). In addition, there are already data suggesting that intrathecal curcumin may reduce pain by inhibiting the activation of astrocytes and microglia and the production of corresponding glial-derived inflammatory mediators such as IL-1β, MCP-1, and MIP-1α in the spinal cord(Zhang et al., 2016, 2012). It was found that in the rat Dorsal root ganglion (DRG) chronic compression model, the up-regulation of p-ERK1/2 expression in DRG can promote the activation of large and medium-diameter neurons, and the blocking of ERK1/2 signaling pathway can inhibit neuronal activation and reduce the continuous high response of DRG, effectively alleviating the mechanical pain sensitivity of rats, indicating that ERK1/2 in peripheral nerves promotes information exchange between neurons and glial cells(Ji et al., 2009). ERK1/2 is an upstream regulatory kinase of CDK5, and the possible mechanism of CDK5 involvement in inflammatory pain is that p-ERK1/2 upregulates the expression of p35 by phosphorylating early growth response factor 1 in the nucleus, while p35 binds to CDK5 to stimulate the enzymatic activity of CDK5, inhibits the outward current of potassium ion A ions by regulating the expression of potassium channel Kv4.2 in dorsal horn neurons of the spinal cord, lowers the excitatory threshold of neurons, and changes synaptic plasticity(Pareek et al., 2013; Qu et al., 2016). Studies have shown that the activation of the spinal cord Cdk5/p35 signaling pathway can regulate the transcription of DNA by phosphorylating a variety of transcription factors, and then promote the expression of a variety of kinase receptors, ion channel receptors or inflammatory factors, which can increase neuronal excitability or decrease the excitability threshold, and change synaptic plasticity. At present, studies on the role of the CDK5/P53 signaling pathway in glial cells in the development of pain have not been reported. Therefore, we tried to explore the interaction between glial cells and the cdk5/p53 signaling pathway in the process of pain development, and explore whether curcumin exerts an analgesic effect by interfering with the CDK5/P35 signaling pathway in glial cells, and then find therapeutic drugs for NP by using the analgesic drug curcumin.

Currently, the mechanism by which curcumin modifies microglial polarization and neuroinflammation during neuropathic pain is unclear(Gao et al., 2019). Given the multifaceted effects of curcumin, we hypothesize that it may exert its analgesic effects by inhibiting p35/CDK5 expression within microglia. By doing so, curcumin could potentially dampen nerve inflammation in critical regions such as the dorsal horn of the spinal cord and the dorsal root ganglion, ultimately leading to a reduction in neuropathic pain symptoms. Further investigation is warranted to validate this hypothesis and elucidate the underlying mechanisms.

## 2 Materials and methods

### 2.1 Animals

The experimental animals used in this study were male Sprague-Dawley (SD) rats, weighing between 220-250 grams, sourced from the Shanghai Experimental Animal Centre in China. All experimental procedures were approved by the Animal Research and Education Committee of Zhejiang University School of Medicine. The number of animals used in all experiments was kept to a minimum in accordance with guidelines set by the International Association for the Study of Pain Ethical Issues. The rats were maintained in a controlled environment with a 12-hour light/dark cycle and had unrestricted access to food and water throughout the study period. The experiment was divided into two phases.

### 2.2 Induction of NP

After the baseline measurements were established, the rats were randomly allocated to either the sham group or the chronic constriction injury (CCI) group. Under anesthesia with 1% pentobarbital sodium(40mg/kg)(Wu et al., 2020), the left sciatic nerve of each rat was exposed by carefully dissecting the middle thigh without sharp instruments(Zhang et al., 2023). For the CCI group, the sciatic nerve was isolated and then ligated with three separate 4-0 gauge wire, each spaced 1 mm apart. The muscular and dermal tissues were then meticulously sutured closed using 4-0 sutures, layer by layer. In contrast, the sham group underwent visualization of the left sciatic nerve without any ligation. Following the surgical procedures, all rats received a subcutaneous injection of 80,000 units of penicillin to guard against post-operative infections.

### 2.3 Behavioral tests

Before experiments, all rats were acclimated to the climate in a 12 cm × 15 cm × 22 cm plastic box placed on an elevated wire mesh for 30 minutes each day over 3 consecutive days. The behavioral experimenter was unaware of the experimental protocol.

### **2.3.1** Thermal Withdrawal Latency (TWL)

Using the Hargreaves method, the latency of the thermal foot contraction response in rats was assessed at various time points using the XR1102 Smart Thermal Stimulator Pain Apparatus(Holden et al., 2014). The time of quick withdrawal or licking of the paw was taken as the TWL. To avoid burn injuries, stimulation was terminated if no positive response was observed 20 s after initiation. Each rat received three trials with an interval of 10 min, and the average values were recorded as the final TWL for analysis.

### **2.3.2** Mechanical Withdrawal Threshold (MWT)

To evaluate mechanical nociceptive abnormalities, paw contraction thresholds were assessed in response to von Frey fine filament stimulation as described previously(Ma et al., 2016). Briefly, the left plantar surface was stimulated with filaments of increasing stiffness (0.4–26 g) until a quick withdrawal or licking of the paw was noted, and the magnitude of the filaments was recorded as the MWT. The testing was repeated three times with an interval of 5 min, and the average value was considered the final MWT.

### **2.3.3** Acetone-induced cold allodynia test

The acetone test score (ATS) was used to assess cold allodynia, as detailed by Farsi et al. (Farsi et al., 2015; Yuan et al., 2020) Using the same apparatus as for the MWT test. In this test, 100µL of acetone was sprayed onto the left plantar surface, and the response was observed for 20 seconds. Responses were scored on a 4-point scale ranging from no response (0) to prolonged and repetitive withdrawal with flinching/licking (4). The test was repeated three times with 5-minute intervals, and the average score was taken as the final ATS.

### 2.4 Immunofluorescence

After deep anesthesia with pentobarbital sodium, the rats were transcardially perfused with 150 mL of 1 × phosphate buffered saline (PBS) (4 °C), followed by 150 mL of 4% paraformaldehyde (4 °C). The L4-L6 spinal cord tissue and dorsal root ganglia was harvested, fixed with 4% paraformaldehyde for 48 h, and then dehydrated with dehydrated in a series of 20% and 30% sucrose solutions for 3 days at 4 °C. Subsequently, the ACC was transversely cut into slices (30-µm thick). The sections were blocked with 10% sheep serum for 2 h at room temperature and incubated with the following primary antibodies for 48 h at 4 °C: goat-anti-Iba1 (1:100, Abcam), rabbit anti-p35(1:400, Cell Signaling Technology), rabbit anti-CDK5 (1:500, Cell Signaling Technology). The sections were washed with 1 × PBS and incubated with fluorescent secondary antibodies in the dark for 2 h at room temperature. Finally, the sections were examined under a fluorescence microscope (FV3000; Olympus, Tokyo, Japan).

### 2.5 Western blotting

Western blotting was conducted according to established protocols (Chen et al., 2020; Guo et al., 2009). Rats were anesthetized with pentobarbital sodium, decapitated, and their brains were harvested. The L4-L6 spinal cord and dorsal root ganglia were isolated and placed in a centrifuge tube with 500 μL of protein lysis buffer. The supernatant was then transferred to another tube, and total protein content was measured using a BCA kit. The proteins were separated by SDS-PAGE and transferred to a PVDF membrane. Membranes were cut based on protein markers, blocked with 5% skimmed milk for 1 hour, and incubated with primary antibodies (mouse-anti-CDK5 1:200,rabbit-anti-p35 1:500, mouse-anti-Tubulin 1:2000) for 24 hours at 4 °C. After washing with TBST, the membrane was incubated with a secondary antibody for 2 hours at room temperature. Protein bands were visualized using an ECL kit and captured with a gel imaging system. Band density was analyzed with Image Lab software to quantify target protein expression.

### 2.6 Statistical analysis

For data analysis, two-way repeated measure analysis of variance was used, followed by Tukey’s post-hoc test. P <0.05 was considered significant. All data are expressed as Mean ± SEM. SPSS18 was used for analysis (SPSS Inc., Chicago, Illinois, USA).

## 3 Results

### 3.1 Curcumin effectively improves pain sensitization in rats with neuropathic pain

To explore the role of curcumin in pain modulation, we first investigated the regulatory role of curcumin on pain threshold changes. (Fig.1A) No significant difference was observed in the baseline pain behavior between the sham and CCI groups before surgery. Seven days after surgery, the MWT and TWL in the CCI+DMSO group were significantly lower than those in the sham group, alongside a marked increase in acetone-induced cold allodynia scores in the CCI+DMSO group. These results suggest that CCI successfully induced behavioral hyperalgesia. Compared with the CCI+DMSO group, following 4 days of curcumin treatment in the CCI+Cur group, MWT and TWL values were significantly higher and acetone-induced cold allodynia test scores were significantly lower. These results suggest that curcumin plays an important role in the regulation of neuropathic pain. (Fig.1B-D) In addition, no differences were observed in the MWT, TWL, and acetone cold hyperalgesia test scores of the contralateral hind paw at each time point in each group. (Fig.1E-G)

**Figure 1.**
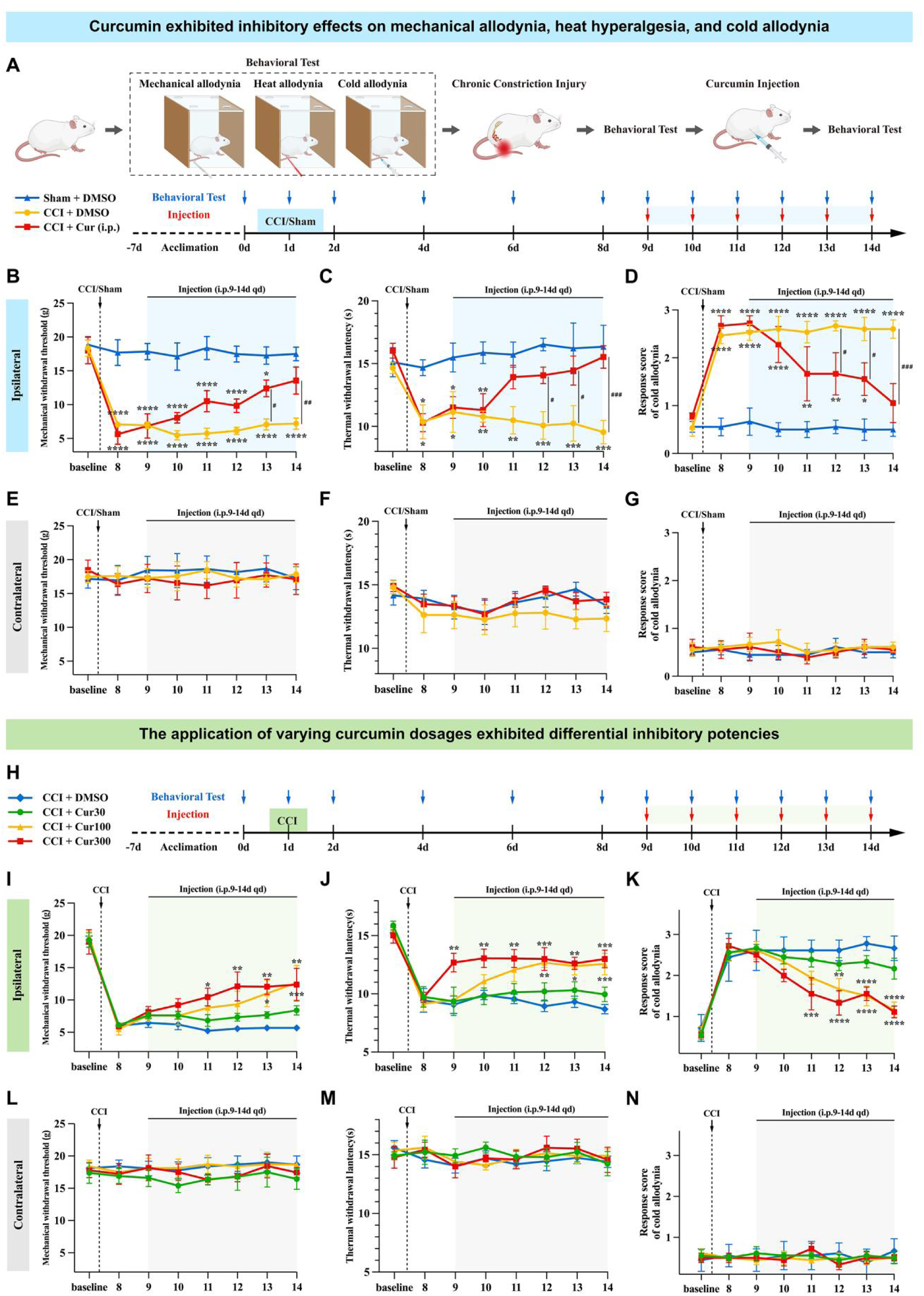
Analgesic effects of curcumin at different doses on mechanical allodynia, heat hyperalgesia and cold allodynia in neuropathic pain rats. **(A)** Experimental protocol for CCI or Sham surgery, behavioral test and injection of curcumin in the Sham+DMSO group, the CCI+DMSO group and the CCI+Cur group. **(B, E)** Results showed the effects of i.p. injection of curcumin on the ipsilateral and contralateral MWT. **(C, F)** Results showed the effects of i.p. injection of curcumin on the ipsilateral and contralateral TWL. **(D, G)** Results showed the effects of i.p. injection of curcumin on the ipsilateral and contralateral acetone-induced cold allodynia. **(H)** Experimental protocol for CCI surgery, behavioral test and different doses of injection of curcumin in the CCI+DMSO group, the CCI+Cur30 group, the CCI+Cur100 group and the CCI+Cur300 group. **(I, L)** Results showed the effects of i.p. injection of curcumin with different doses on the ipsilateral and contralateral MWT. **(J, M)** Results showed the effects of i.p. injection of curcumin with different doses on the ipsilateral and contralateral WTL. **(K, N)** Results showed the effects of i.p. injection of curcumin with different doses on the ipsilateral and contralateral acetone-induced cold allodynia. \*\*\*\**P* < 0.0001, \*\*\**P* < 0.001, \*\**P* < 0.01, \**P* < 0.05; #*P* < 0.05; ##*P* < 0.01, ###*P* < 0.001; n=6/group CCI: chronic compression injury; Cur: curcumin; i.p.: intraperitoneal injection; MWT: mechanical withdrawal threshold; TWL: thermal withdrawal latency;

### 3.2 Curcumin dose-dependently exhibits different inhibitory potencies for neuropathic pain hyperalgesia

Based on the above results, we have identified the huge role played by curcumin in regulating pain threshold changes. To further explore the role of curcumin in neuropathic pain, we injected different doses of curcumin continuously from day 9 to day 14 after CCI. (Fig.1H) Behavioral analysis revealed no significant differences in MWT, TWL, or acetone-induced cold allodynia scores between the CCI+DMSO and CCI+Cur30 groups. However, after 4 days of curcumin treatment in the CCI+Cur100 group, MWT and TWL increased significantly, while acetone-induced cold allodynia scores decreased. Similarly, a 3-day treatment with 300 mg/kg of curcumin in the CCI+Cur300 group elevated MWT and TWL, indicating reduced sensitivity to mechanical and thermal stimuli, respectively, and alleviated cold hyperalgesia. (Fig. 1I-K) The results showed that compared with that in the CCI+DMSO group, curcumin not only plays a crucial role in modulating neuropathic pain but also exhibits a dose-dependent regulation of pain threshold changes. Similarly, no differences were found in these parameters for the contralateral hind paw in any group at any time point, indicating that curcumin has no effect on normal tissues. (Fig. 1L-N)

### 3.3 Curcumin reverses the activation of p35/CDK5 pain-related signaling factors in the spinal cord and appears to be dose-dependent

To determine whether curcumin exerts an influence within the dorsal horn of the spinal cord and to identify the mediators involved, we conducted further experiments to analyze. (Fig.2A) Consistent with behavioral hyperalgesia observed after CCI, western blotting results showed overexpression of p35/CDK5 protein compared to the sham group, while curcumin use significantly reversed this effect. These results indicated that the curcumin alleviates neuropathic pain by targeting p35/CDK5 protein in the ipsilateral spinal cord. Western blot results showed that CCI surgery and intraperitoneal injection of curcumin had no significant effect on the expression of p35/CDK5 protein in the dorsal horn of the ipsilateral spinal cord. (Fig.2B-I) We speculate that by inhibiting the increase in p35/CDK5 expression in the spinal cord, curcumin may disrupt the signaling cascades that contribute to pain facilitation and neuroinflammation. In addition, the expression of p35/CDK5 protein differed under different doses of curcumin. Compared with the CCI+DMSO group, there was no significant difference in p35/CDK5 expression in the CCI+Cur30 group. The expression of p35/CDK5 protein in the spinal cord group of rats treated with curcumin (100 or 300 mg/kg) was significantly reduced. (Fig.2K-R) Similarly, there was no significant change in the p35/CDK5 protein in the dorsal horn of the contralateral spinal cord, which was consistent with the behavioral results. These results suggest that p35/CDK5 protein is an important target for curcumin to regulate neuropathic pain, and it reverses the pain signal expression of NP in a dose-dependent manner.

**Figure 2.**
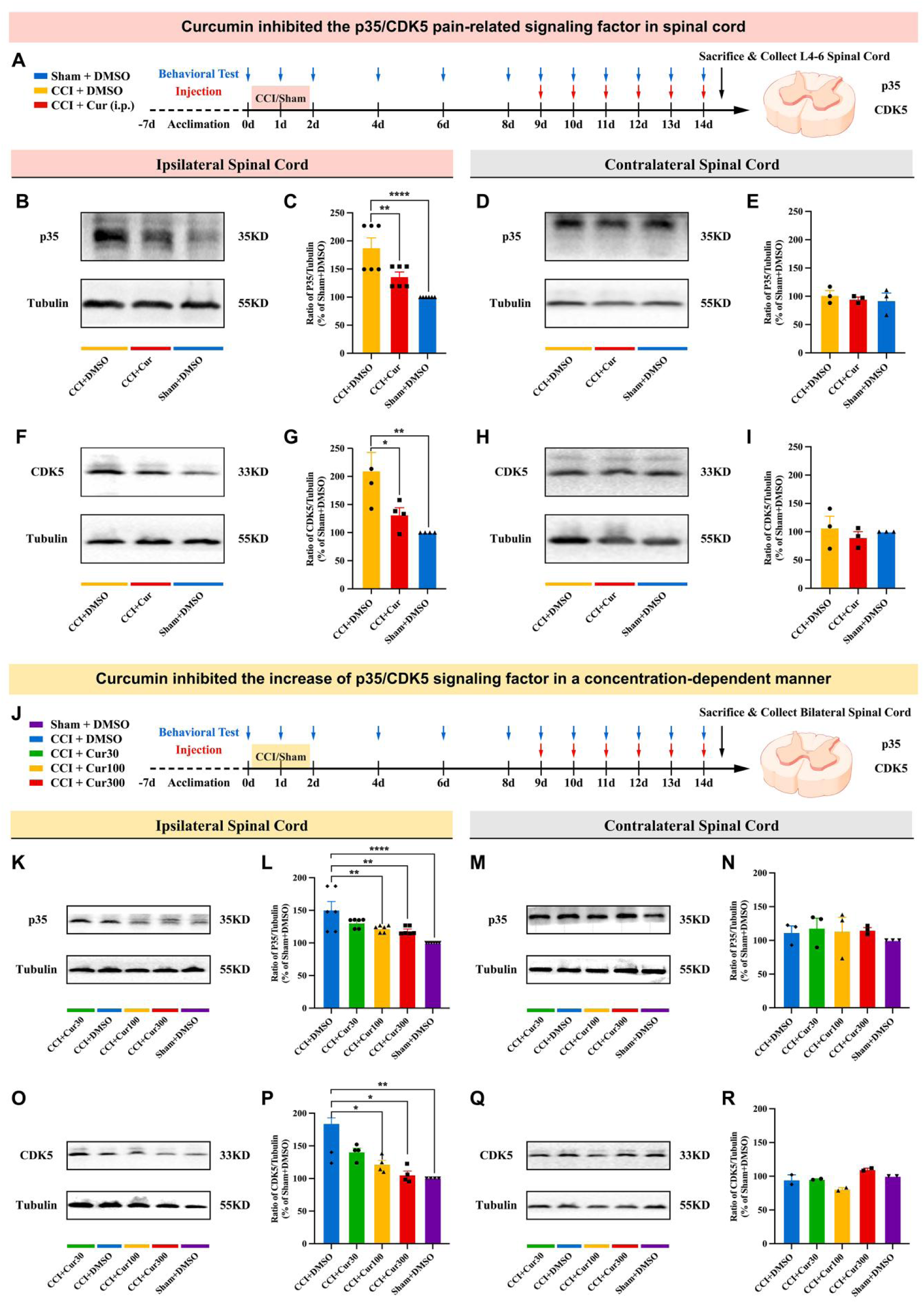
Curcumin influences the expression of p35/CDK5 in the bilateral spinal cord caused by CCI model. **(A)** Experimental protocol for CCI or Sham surgery, behavioral test, injection of curcumin and collection of bilateral spinal cord in the Sham+DMSO group, the CCI+DMSO group and the CCI+Cur group. **(B, D)** Representative images showed WB analysis of p35 expression in ipsilateral and contralateral spinal cord. **(C, E)** Results showed the effects of i.p. injection of curcumin on the p35 expression in the ipsilateral and contralateral spinal cord. **(F, H)** Representative images showed WB analysis of CDK5 expression in ipsilateral and contralateral spinal cord. **(G, I)** Results showed the effects of i.p. injection of curcumin on the CDK5 expression in the ipsilateral and contralateral spinal cord. **(J)** Experimental protocol for CCI or Sham surgery, behavioral test, injection of curcumin and collection of bilateral spinal cord in the CCI+DMSO group, the CCI+Cur30 group, the CCI+Cur100 group and the CCI+Cur300 group. **(K, M)** Representative images showed WB analysis of p35 expression in ipsilateral and contralateral spinal cord. **(L, N)** Results showed the effects of i.p. injection of curcumin with different doses on the p35 expression in the ipsilateral and contralateral spinal cord. **(O, Q)** Representative images showed WB analysis of CDK5 expression in ipsilateral and contralateral spinal cord. **(P, R)** Results showed the effects of i.p. injection of curcumin with different doses on the CDK5 expression in the ipsilateral and contralateral spinal cord. \*\*\*\**P* < 0.0001, \*\**P* < 0.01, \**P* < 0.05, n=6/group ; CCI: chronic compression injury; Cur: curcumin; WB: western blotting; i.p.: intraperitoneal injection; CDK5: Cyclin-dependent kinase 5

### 3.4 Curcumin reverses the involvement of spinal dorsal horn microglia in the regulation of neuropathic pain and is closely related to p35/CDK5 protein

The results of the above experiments have confirmed that p35/CDK5 is involved in the modulation of spinal cord pain signals, but the role of glial cells in synaptic plasticity regulation during chronic pain maintenance cannot be ignored, so we introduced Iba-1 to determine whether microglia are involved in central sensitization. The results of immunofluorescence co-staining showed that p35/CDK5 and microglia marker Iba-1 were co-expressed in the dorsal horn of the spinal cord of SD rats after CCI, suggesting that neuropathic pain was highly correlated with microglial activation, and intraperitoneal injection of curcumin could effectively reverse the co-expression of microglial marker Iba-1 and p35/CDK5, which reflected the direct inhibitory effect of curcumin on microglial activation, and suggested curcumin may exert its analgesic effects, at least partially, by modulating CDK5 signaling and microglial activation in the spinal cord. (Fig.3A-E) Overall, our results suggest that the expression of p35 and CDK5 is intimately associated with microglial activation and that curcumin represents a potential therapeutic agent for modulating this process in the context of pain and neuroinflammation.

**Figure 3.**
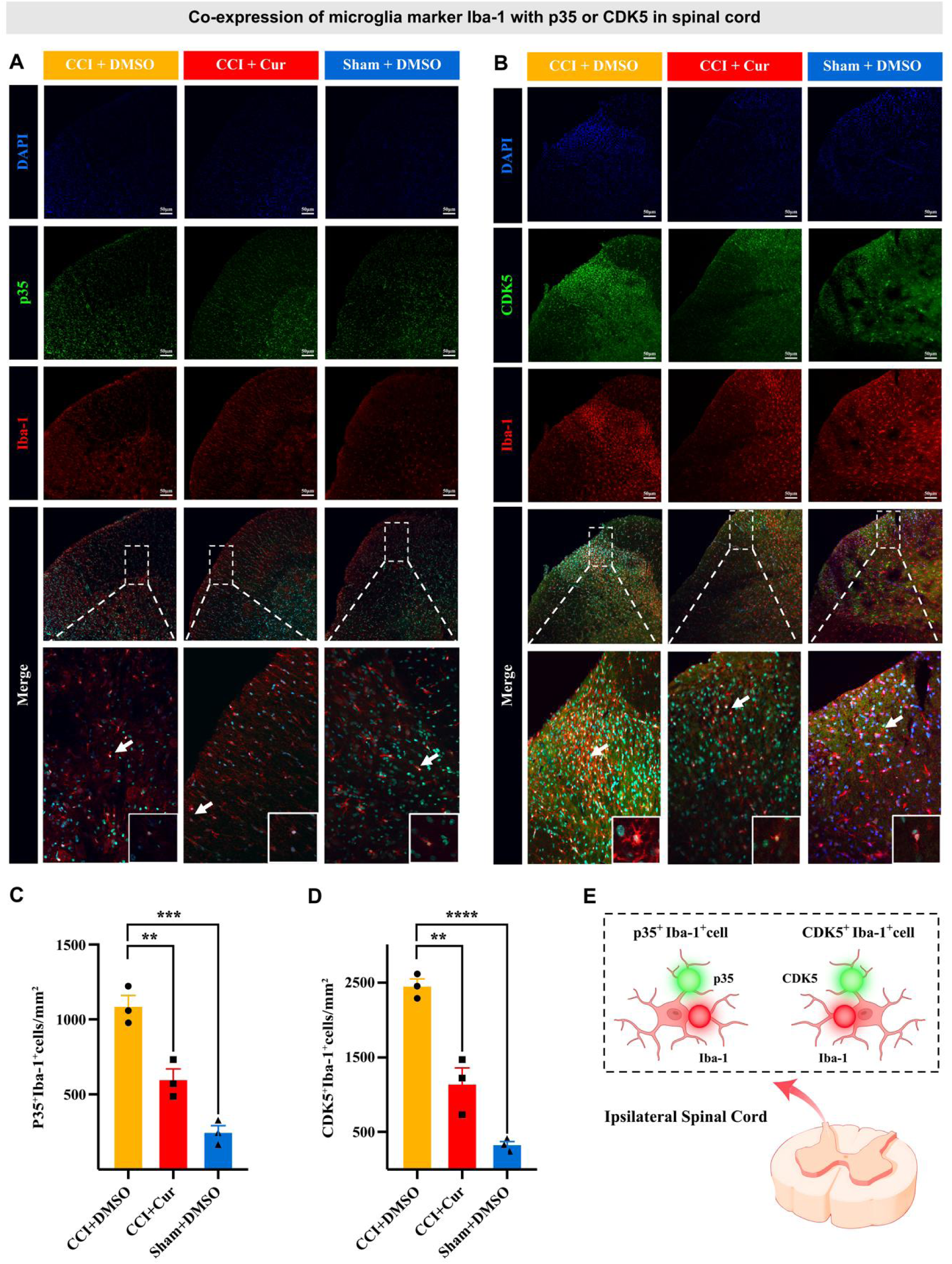
Double immunofluorescence of p35 or CDK5, and microglial marker Iba-1 in spinal cord of CCI rats. **(A)** Representative images of staining for p35, Iba-1, and DAPI in the ipsilateral spinal cord in the Sham+DMSO group, the CCI+DMSO group and the CCI+Cur group.. **(B)** Representative images of staining for CDK5, Iba-1, and DAPI in the ipsilateral spinal cord in the Sham+DMSO group, the CCI+DMSO group and the CCI+Cur group. **(C)** Results showed the effects of i.p. injection of curcumin on the co-expression of p35 and Iba-1 in the ipsilateral spinal cord. **(D)** Results showed the effects of i.p. injection of curcumin on the co-expression of CDK5 and Iba-1 in the ipsilateral spinal cord. **(E)** Schematic diagram of the co-expression of microglia marker Iba-1 and p35 or CDK5 in the ipsilateral spinal cord. \*\*\*\**P* < 0.0001, \*\*\**P* < 0.001, \*\**P* < 0.01,n=6/group ; CCI: chronic compression injury; Cur: curcumin; i.p.: intraperitoneal injection; CDK5: Cyclin-dependent kinase 5

### 3.5 Curcumin effectively suppressed the p35/CDK5 pain-related signaling pathway in the DRG, exhibiting a concentration-dependent inhibitory effect on the upregulation of p35/CDK5 signaling factors

The dorsal root ganglia acts as a pre-sequence signal station for pain transmission to the dorsal horn of the spinal cord, and whether curcumin is affected in the inhibition of pain conduction remains to be further clarified. The results of western blot analysis showed that the expression of p35/CDK5 protein in the CCI+DMSO group was significantly increased compared with the Sham+DMSO group. Compared with the CCI+DMSO group, the expression of p35/CDK5 treated with curcumin was significantly reduced in the dorsal root ganglia of rats. (Fig.4B-I) On the one hand, it is clear that the p35/CDK5 protein in the dorsal root ganglia is also heavily activated, that is, the dorsal root ganglia is involved in regulating the formation of pain sensitization, and on the other hand, it also proves that curcumin not only plays an inhibitory role in the dorsal horn of the spinal cord, but also plays an important role in the dorsal root ganglia.

**Figure 4.**
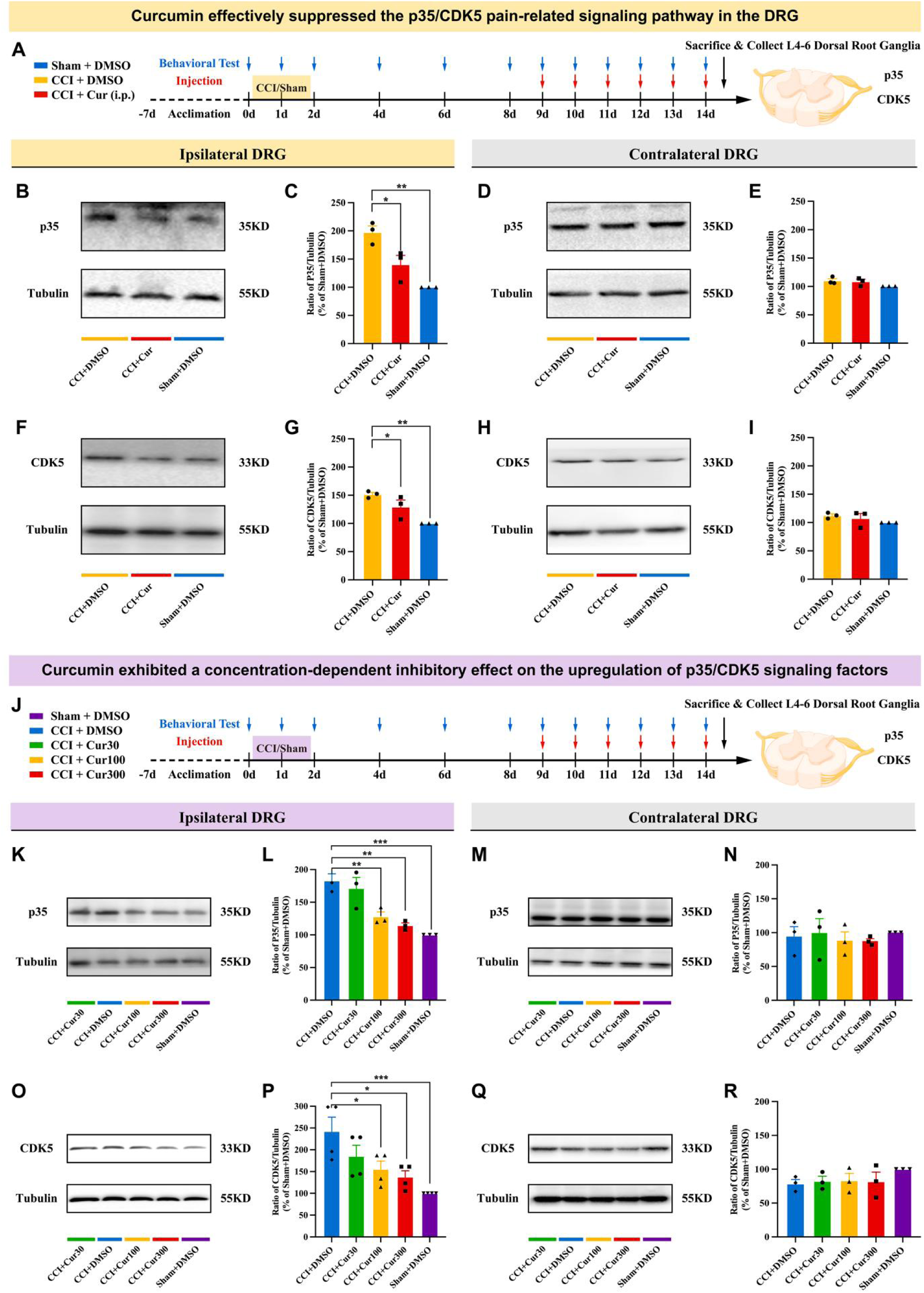
Curcumin influences the expression of p35/CDK5 in the bilateral lumbar 4-6th dorsal root ganglia caused by CCI model. **(A)** Experimental protocol for CCI or Sham surgery, behavioral test, injection of curcumin and collection of bilateral L4-6 DRG in the Sham+DMSO group, the CCI+DMSO group and the CCI+Cur group. **(B, D)** Representative images showed WB analysis of p35 expression in ipsilateral and contralateral DRG. **(C, E)** Results showed the effects of i.p. injection of curcumin on the p35 expression in the ipsilateral and contralateral DRG. **(F, H)** Representative images showed WB analysis of CDK5 expression in ipsilateral and contralateral DRG. **(G, I)** Results showed the effects of i.p. injection of curcumin on the CDK5 expression in the ipsilateral and contralateral DRG. **(J)** Experimental protocol for CCI or Sham surgery, behavioral test, injection of curcumin and collection of bilateral L4-6 DRG in the CCI+DMSO group, the CCI+Cur30 group, the CCI+Cur100 group and the CCI+Cur300 group. **(K, M)** Representative images showed WB analysis of p35 expression in ipsilateral and contralateral DRG. **(L, N)** Results showed the effects of i.p. injection of curcumin with different doses on the p35 expression in the ipsilateral and contralateral DRG. **(O, Q)** Representative images showed WB analysis of CDK5 expression in ipsilateral and contralateral DRG. **(P, R)** Results showed the effects of i.p. injection of curcumin with different doses on the CDK5 expression in the ipsilateral and contralateral DRG. \*\*\**P* < 0.001, \*\**P* < 0.01, \**P* < 0.05, n=6/group; CCI: chronic compression injury; DRG: dorsal root ganglia; Cur: curcumin; WB: western blotting; i.p.: intraperitoneal injection; CDK5: Cyclin-dependent kinase 5

Additionally, p35/CDK5 expression varied at different doses of curcumin. In comparison to the CCI+DMSO group, the CCI+Cur30 group exhibited no notable difference in p35/CDK5 expression. The expression of p35/CDK5 protein treated with curcumin (100 or 300 mg/kg) was also significantly reduced (Fig.4K-R). These findings demonstrate that curcumin inhibits the upregulation of p35/CDK5 pain-related factors in a concentration-dependent manner within the lumbar 4-6th dorsal root ganglia. This inhibitory effect aligns with similar observations made in the spinal cord, suggesting a consistent mechanism of action across these neural tissues.

### 3.6 Curcumin reverses dorsal root ganglia microglia involved in the regulation of neuropathic pain and is closely related to p35/CDK5 protein

Now that the role of dorsal root ganglia in curcumin in improving pain threshold has been clarified, a large number of immune cells are also gathered in the dorsal root ganglia, and similarly, we also want to explore the changes of microglia in pain function. In each experimental group, co-expression of Iba-1 with p35 and CDK5 markers was detected in lumbar DRGs spanning segments 4 to 6. P35 was co-expressed with the microglia marker Iba-1 (Figure 5A). Similarly, the co-expression of CDK5 with Iba-1 further supports the involvement of this kinase in microglial function (Figure 5B). Curcumin intervention attenuated the co-expression of Iba-1, p35, and CDK5 compared to the CCI+DMSO group (Fig. 5C-D). This observation is consistent with previous findings in the spinal cord.

**Figure 5.**
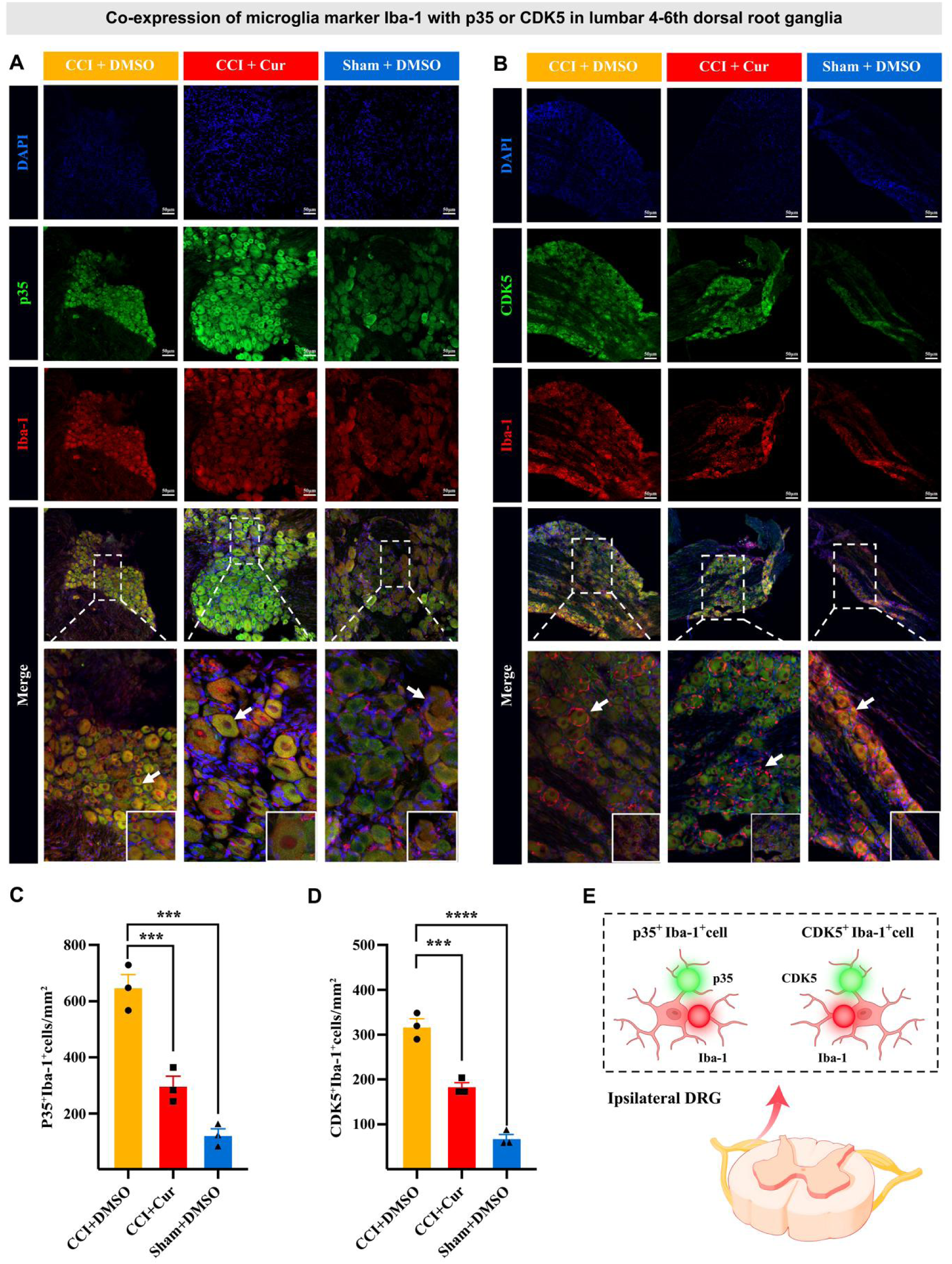
Double immunofluorescence of p35 or CDK5, and microglial marker Iba-1 in lumbar 4-6th dorsal root ganglia of CCI rats. **(A)** Representative images of staining for p35, Iba-1, and DAPI in the ipsilateral L4-6 DRG in the Sham+DMSO group, the CCI+DMSO group and the CCI+Cur group. **(B)** Representative images of staining for CDK5, Iba-1, and DAPI in the ipsilateral DRG in the Sham+DMSO group, the CCI+DMSO group and the CCI+Cur group. **(C)** Results showed the effects of i.p. injection of curcumin on the co-expression of p35 and Iba-1 in the ipsilateral DRG. **(D)** Results showed the effects of i.p. injection of curcumin on the co-expression of CDK5 and Iba-1 in the ipsilateral DRG. **(E)** Schematic diagram of the co-expression of microglia marker Iba-1 and p35 or CDK5 in the ipsilateral DRG. \*\*\*\**P* < 0.0001, \*\*\**P* < 0.001,n=6/group ; DRG: dorsal root ganglia; CCI: chronic compression injury; Cur: curcumin; i.p.: intraperitoneal injection; CDK5: Cyclin-dependent kinase 5

Based on the above experimental results, we confirmed that CCI-induced sciatic nerve injury can lead to neuropathic pain, increase CDK5 and p35 protein activity, and activate microglia in the dorsal horn and dorsal root ganglia of the spinal cord, and clarify the interaction between neurons and immune cells in the regulation of neuropathic pain. Curcumin, as a traditional Chinese medicine preparation, has a significant analgesic effect on CCI-induced neuropathic pain by inhibiting p35/CDK5 activity and microglial activation. Curcumin may be an effective analgesic for the treatment of neuropathic pain; In addition, the activation of microglia in the late stages of nerve injury may be a good target for the treatment of NPs that have already formed.

## 4 Discussion

Neuropathic pain (NP) is a complex and challenging clinical condition that remains a significant burden for patients and healthcare systems worldwide. NP arises from damage or dysfunction in the somatosensory system, manifesting as spontaneous pain, hyperalgesia, allodynia, hypoesthesia, and emotional disturbances. Recent discoveries have underscored the pivotal role of microglia in the central nervous system, particularly at the spinal cord level, where they modulate neuronal function through direct connections, synaptic interactions, or the release of neurotransmitters. Furthermore, microglia contribute to NP pathology by facilitating long-term potentiation (LTP), playing a crucial role in the onset and progression of NP and offering potential therapeutic targets. In recent years, research has witnessed a surge in the potential therapeutic effects of curcumin in treating this debilitating condition. However, the precise mechanisms underlying curcumin’s treatment of neuropathic pain remain elusive. Therefore, we conducted a series of studies aimed at elucidating how curcumin influences microglial activation in the spinal cord and dorsal root ganglia (DRG) following chronic constriction injury (CCI) of the rat sciatic nerve, particularly through the p35/CDK5 signaling pathway.

To further investigate the effects of curcumin on microglial activation in the spinal cord and DRG of rats with CCI via the p35/CDK5 signaling pathway, we established a neuropathic pain model through chronic constriction injury (CCI) of the sciatic nerve and administered varying doses of curcumin to form curcumin-treated groups. Our findings demonstrate that curcumin exhibits potent analgesic properties by alleviating mechanical pain, thermal nociception, and cold nociceptive hyperalgesia in CCI rats. Our behavioral test results reveal that curcumin treatment significantly improved mechanical withdrawal threshold (MWT) and thermal withdrawal latency (TWL) in CCI rats compared to the vehicle-treated group. Furthermore, curcumin reduced cold allodynia, as evidenced by lower acetone-induced cold allodynia test scores. These results are consistent with previous studies that have reported curcumin’s analgesic effects in various pain models(Eke-Okoro et al., 2018). However, our study specifically highlights curcumin’s efficacy in alleviating neuropathic pain, which is known to be particularly challenging to treat.

The dose-response relationship observed in our study is crucial for the clinical translation of curcumin as a therapeutic agent for neuropathic pain. We found that curcumin at doses of 100 mg/kg and 300 mg/kg exhibited significant analgesic effects, with the higher dose showing a quicker onset of action. Conversely, a lower dose of 30 mg/kg did not significantly alleviate pain symptoms. These findings inform dosing strategies that balance efficacy and safety, which are essential considerations in the development of new therapeutic interventions.

Neuropathic pain is strongly associated with microglial activation in the spinal cord and DRG (An et al., 2021; Yu et al., 2019). Microglia are resident macrophages in the brain and spinal cord of the central nervous system, and there is substantial evidence to support the role of microglia in neuropathic pain processes(Grace et al., 2014; Ji et al., 2013). One of the major characteristics of microglia is that they are highly sensitive to environmental surveillance, and even if there are slight pathological changes in the central nervous system, they will activate rapidly to exert immune function. Our immunofluorescence and western blotting results showed that curcumin potently inhibited microglial activation, manifested by a decrease in the expression of the microglia marker Iba-1. In addition, we also found that curcumin down-regulated the expression of p35 and CDK5, which are key components of the p35/CDK5 signaling pathway. This finding is particularly important because CDK5 plays a key role in pain signaling and neuroinflammation when activated by its cofactor p35 (Chapman et al., 2019; Zeb et al., 2019). Our data suggest that curcumin’s analgesic effects may be mediated through inhibiting p35/CDK5 signaling in activated microglia. This inhibition likely reduces neuroinflammation, which is a key contributor to neuropathic pain development and maintenance. Consistent with this hypothesis, we observed that curcumin treatment reduced the co-expression of Iba-1 with p35 and CDK5 in both the spinal cord and DRG, further supporting its ability to modulate microglial activation and signaling. These results suggest that microglia play a key role in the early formation of neuropathic pain. Long-term pain and sustained microglial activation can lead to structural and functional alterations in the pain pathway, i.e., alterations in neuroplasticity. Studies have shown that CDK5 binding stimulates the enzymatic activity of CDK5, and activated CDK5 regulates the expression of Kv4.2 potassium channels in dorsal horn neurons of the spinal cord, inhibits the outward current of type A potassium ions, lowers the excitability threshold of neurons, and alters synaptic plasticity(Ji et al., 2009; Pareek et al., 2013). Therefore, in the next research, we will pay more attention to the complex and critical role of microglia in the occurrence, development and maintenance of neuropathic pain through these pathways, and explore the mechanism of curcumin on NP in the process of chronic pain

Our data suggest that curcumin’s analgesic effects may be mediated through inhibiting p35/CDK5 signaling in activated microglia. This inhibition likely reduces neuroinflammation, which is a key contributor to neuropathic pain development and maintenance. Consistent with this hypothesis, we observed that curcumin treatment reduced the co-expression of Iba-1 with p35 and CDK5 in both the spinal cord and DRG, further supporting its ability to modulate microglial activation and signaling.

However, although our study provides strong evidence for the analgesic effect of curcumin and its regulatory effect on the p35/CDK5 signaling pathway, the exact molecular mechanisms of these effects remain to be fully elucidated. Curcumin may inhibit other signaling pathways or act on multiple components of the pain cascade. A large number of studies have confirmed that MAPK family members play an important role in the occurrence and maintenance of pain and can participate in the modulation of spinal pain signals by promoting glial cell activation(Ji et al., 2009; Norsted Gregory et al., 2013; Tawfik et al., 2007). At the same time, as a regulator of glial cell activation, PPF may have some analgesic effect by inhibiting glial cell MAPK kinases(Liu et al., 2020; Norsted Gregory et al., 2013; Wu et al., 2013), thereby exerting its observed therapeutic effects. Future studies are needed to explore these potential mechanisms and identify other curcumin targets related to neuropathic pain.

Overall, we found that curcumin downregulated p35 and CDK5, reversing pain sensitization in rats with the chronic constriction injury (CCI) model, indicating its significant potential as a therapeutic option for neuropathic pain. However, studies have also shown that curcumin may inhibit CX3CR1 expression by activating NF-κB p65 in the spinal dorsal horn and dorsal root ganglion (DRG), thereby alleviating neuropathic pain in CCI(Liu et al., 2020). Furthermore, hormonal responses, such as those involving corticosterone (CORT), and curcumin can reduce mechanical and thermal hyperalgesia by inhibiting serum cortisol concentrations and decreasing the expression of 11β-hydroxysteroid dehydrogenase type 1 (11bHSD1) in superficial spinal dorsal horn and DRG neurons(Cao et al., 2014; Di et al., 2014). Results from another study demonstrated that curcumin significantly reduced calcium accumulation in the sciatic nerve, decreased nitric oxide (NO) and lipid peroxidation (LPO), and increased endogenous antioxidant enzymes in vincristine-induced neuropathic pain(Babu et al., 2015). In summary, as a natural compound extracted from turmeric, curcumin possesses advantages such as low toxicity, good bioavailability, and a wide therapeutic window that traditional drugs lack(Li et al., 2022). Its effects are not exerted through a single target or pathway; rather, its various actions, including anti-nociceptive, calcium inhibitory, and antioxidant effects, are achieved through the combined influence on different protein molecules. It is important to focus not only on the effects of curcumin on different pathways but also on the interactions between these pathways. The drug interactions when used in combination with other medications cannot be ignored. In future research, it is necessary to conduct clinical trials to evaluate the efficacy and safety of curcumin in patients with neuropathic pain, with special attention to optimizing dosing regimens and exploring combination therapies.

In conclusion, our study demonstrates that curcumin alleviates neuropathic pain in rats by modulating microglia activation and inhibiting the p35/CDK5 signaling pathway. These findings contribute to our understanding of curcumin’s analgesic mechanisms and highlight its potential as a novel therapeutic agent for neuropathic pain management.

## 5 Conclusion

Curcumin attenuates ipsilateral mechanical pain, thermal nociception and cold nociceptive hyperalgesia in CCI rats. According to the analysis of the present study, curcumin may alleviate neuropathic pain in rats by modulating microglia in the spinal cord and dorsal root ganglia through the p35/CDK5 signaling pathway.

## Authors’ contributions

Liang Yu, lei Zhang YongXing Yao and HongHai Zhang developed the concept of the study and designed the study;Fei Tong, li Zhao, Yue Yang ZhuoYue Zhang, Lu Liu, YaXuan Wu performed the experiments and analyzed the data.XuanYi Di, ZiWen Zhang, XiaoXia Xu, YuanLi Zhang Yue Shen helped with the analysis of the experiments. All authors reviewed and approved the final manuscript.

## Acknowledgments

Thanks to all those who have helped us.

## Data Availability

The data supporting the findings of this study are available within the article. Data will be made available upon reasonable request.

## Declarations

### Ethics Approval

All procedures were in accordance with the National Institutes of Health Guidelines for the Care and Use of Laboratory Animals and were approved by the Animal Advisory Committee of Zhejiang University.

### Consent to Participate

Not applicable.

### Consent for Publication

Not applicable.

## References

[1] An, Q., Sun, C., Li, R., Chen, S., Gu, X., An, S., Wang, Z., 2021. Calcitonin gene-related peptide regulates spinal microglial activation through the histone H3 lysine 27 trimethylation via enhancer of zeste homolog-2 in rats with neuropathic pain. Journal of Neuroinflammation 18, 117. 10.1186/s12974-021-02168-1

[2] Babu, A., Prasanth, K.G., Balaji, B., 2015. Effect of curcumin in mice model of vincristine-induced neuropathy. Pharm Biol 53, 838–848. 10.3109/13880209.2014.943247

[3] Bardin, L., Tarayre, J.P., Malfetes, N., Koek, W., Colpaert, F.C., 2003. Profound, non-opioid analgesia produced by the high-efficacy 5-HT(1A) agonist F 13640 in the formalin model of tonic nociceptive pain. Pharmacology 67, 182–194. 10.1159/000068404

[4] Baron, R., Binder, A., Wasner, G., 2010. Neuropathic pain: diagnosis, pathophysiological mechanisms, and treatment. Lancet Neurol 9, 807–819. 10.1016/S1474-4422(10)70143-5

[5] Bielopolski, N., Amin, H., Apostolopoulou, A.A., Rozenfeld, E., Lerner, H., Huetteroth, W., Lin, A.C., Parnas, M., 2019. Inhibitory muscarinic acetylcholine receptors enhance aversive olfactory learning in adult Drosophila. Elife 8, e48264. 10.7554/eLife.48264

[6] Cao, H., Zheng, J.-W., Li, J.-J., Meng, B., Li, J., Ge, R.-S., 2014. Effects of curcumin on pain threshold and on the expression of nuclear factor κ B and CX3C receptor 1 after sciatic nerve chronic constrictive injury in rats. Chin J Integr Med 20, 850–856. 10.1007/s11655-013-1549-9

[7] Chang, C.-W., Hsiao, Y.-T., Jackson, M.B., 2021. Synaptophysin Regulates Fusion Pores and Exocytosis Mode in Chromaffin Cells. J Neurosci 41, 3563–3578. 10.1523/JNEUROSCI.2833-20.2021

[8] Chapman, D.E., Reddy, B.J.N., Huy, B., Bovyn, M.J., Cruz, S.J.S., Al-Shammari, Z.M., Han, H., Wang, W., Smith, D.S., Gross, S.P., 2019. Regulation of in vivo dynein force production by CDK5 and 14-3-3ε and KIAA0528. Nat Commun 10, 228. 10.1038/s41467-018-08110-z

[9] Chen, J., Harding, S.M., Natesan, R., Tian, L., Benci, J.L., Li, W., Minn, A.J., Asangani, I.A., Greenberg, R.A., 2020. Cell Cycle Checkpoints Cooperate to Suppress DNA- and RNA-Associated Molecular Pattern Recognition and Anti-Tumor Immune Responses. Cell Rep 32, 108080. 10.1016/j.celrep.2020.108080

[10] Cohen, S.P., Mao, J., 2014. Neuropathic pain: mechanisms and their clinical implications. BMJ 348, f7656–f7656. 10.1136/bmj.f7656

[11] Cohen, S.P., Vase, L., Hooten, W.M., 2021. Chronic pain: an update on burden, best practices, and new advances. The Lancet 397, 2082–2097. 10.1016/S0140-6736(21)00393-7

[12] Di, Y.X., Hong, C., Jun, L., Renshan, G., Qinquan, L., 2014. Curcumin attenuates mechanical and thermal hyperalgesia in chronic constrictive injury model of neuropathic pain. Pain Ther 3, 59–69. 10.1007/s40122-014-0024-4

[13] Eke-Okoro, U.J., Raffa, R.B., Pergolizzi, J.V., Breve, F., Taylor, R., NEMA Research Group, 2018. Curcumin in turmeric: Basic and clinical evidence for a potential role in analgesia. J Clin Pharm Ther 43, 460–466. 10.1111/jcpt.12703

[14] Farsi, L., Afshari, K., Keshavarz, M., NaghibZadeh, M., Memari, F., Norouzi-Javidan, A., 2015. Postinjury treatment with magnesium sulfate attenuates neuropathic pains following spinal cord injury in male rats. Behav Pharmacol 26, 315–320. 10.1097/FBP.0000000000000103

[15] Gao, X., Han, Z., Huang, C., Lei, H., Li, G., Chen, L., Feng, D., Zhou, Z., Shi, Q., Cheng, L., Zhou, X., 2022. An anti-inflammatory and neuroprotective biomimetic nanoplatform for repairing spinal cord injury. Bioact Mater 18, 569–582. 10.1016/j.bioactmat.2022.05.026

[16] Gao, Y., Zhuang, Z., Lu, Y., Tao, T., Zhou, Y., Liu, G., Wang, H., Zhang, D., Wu, L., Dai, H., Li, W., Hang, C., 2019. Curcumin Mitigates Neuro-Inflammation by Modulating Microglia Polarization Through Inhibiting TLR4 Axis Signaling Pathway Following Experimental Subarachnoid Hemorrhage. Front Neurosci 13, 1223. 10.3389/fnins.2019.01223

[17] Grace, P.M., Hutchinson, M.R., Maier, S.F., Watkins, L.R., 2014. Pathological pain and the neuroimmune interface. Nat Rev Immunol 14, 217–231. 10.1038/nri3621

[18] Guo, D., Prins, R.M., Dang, J., Kuga, D., Iwanami, A., Soto, H., Lin, K.Y., Huang, T.T., Akhavan, D., Hock, M.B., Zhu, S., Kofman, A.A., Bensinger, S.J., Yong, W.H., Vinters, H.V., Horvath, S., Watson, A.D., Kuhn, J.G., Robins, H.I., Mehta, M.P., Wen, P.Y., DeAngelis, L.M., Prados, M.D., Mellinghoff, I.K., Cloughesy, T.F., Mischel, P.S., 2009. EGFR signaling through an Akt-SREBP-1-dependent, rapamycin-resistant pathway sensitizes glioblastomas to antilipogenic therapy. Sci Signal 2, ra82. 10.1126/scisignal.2000446

[19] Holden, J.E., Wang, E., Moes, J.R., Wagner, M., Maduko, A., Jeong, Y., 2014. Differences in carbachol dose, pain condition, and sex following lateral hypothalamic stimulation. Neuroscience 270, 226–235. 10.1016/j.neuroscience.2014.04.020

[20] Inchingolo, A.D., Inchingolo, A.M., Malcangi, G., Avantario, P., Azzollini, D., Buongiorno, S., Viapiano, F., Campanelli, M., Ciocia, A.M., De Leonardis, N., de Ruvo, E., Ferrara, I., Garofoli, G., Montenegro, V., Netti, A., Palmieri, G., Mancini, A., Patano, A., Piras, F., Marinelli, G., Di Pede, C., Laudadio, C., Rapone, B., Hazballa, D., Corriero, A., Fatone, M.C., Palermo, A., Lorusso, F., Scarano, A., Bordea, I.R., Di Venere, D., Inchingolo, F., Dipalma, G., 2022. Effects of Resveratrol, Curcumin and Quercetin Supplementation on Bone Metabolism-A Systematic Review. Nutrients 14, 3519. 10.3390/nu14173519

[21] Ji, R.-R., Berta, T., Nedergaard, M., 2013. Glia and pain: is chronic pain a gliopathy? Pain 154 Suppl 1, S10–S28. 10.1016/j.pain.2013.06.022

[22] Ji, R.-R., Gereau, R.W., Malcangio, M., Strichartz, G.R., 2009. MAP kinase and pain. Brain Res Rev 60, 135–148. 10.1016/j.brainresrev.2008.12.011

[23] Kong, Y., Shi, W., Zheng, L., Zhang, D., Jiang, X., Liu, B., Xue, W., Kuss, M., Li, Y., Sorgen, P.L., Duan, B., 2023. In situ delivery of a curcumin-loaded dynamic hydrogel for the treatment of chronic peripheral neuropathy. J Control Release 357, 319–332. 10.1016/j.jconrel.2023.04.002

[24] Li, M., Guo, T., Lin, J., Huang, X., Ke, Q., Wu, Y., Fang, C., Hu, C., 2022. Curcumin inhibits the invasion and metastasis of triple negative breast cancer via Hedgehog/Gli1 signaling pathway. Journal of Ethnopharmacology 283, 114689. 10.1016/j.jep.2021.114689

[25] Liu, Z., Yao, X., Jiang, W., Li, W., Zhu, S., Liao, C., Zou, L., Ding, R., Chen, J., 2020. Advanced oxidation protein products induce microglia-mediated neuroinflammation via MAPKs-NF-κB signaling pathway and pyroptosis after secondary spinal cord injury. J Neuroinflammation 17, 90. 10.1186/s12974-020-01751-2

[26] Ma, S.-X., Peterson, R.G., Magee, E.M., Lee, P., Lee, W.-N.P., Li, X.-Y., 2016. Impaired expression of neuronal nitric oxide synthase in the gracile nucleus is involved in neuropathic changes in Zucker Diabetic Fatty rats with and without 2,5-hexanedione intoxication. Neurosci Res 106, 47–54. 10.1016/j.neures.2015.10.007

[27] Norsted Gregory, E., Delaney, A., Abdelmoaty, S., Bas, D.B., Codeluppi, S., Wigerblad, G., Svensson, C.I., 2013. Pentoxifylline and propentofylline prevent proliferation and activation of the mammalian target of rapamycin and mitogen activated protein kinase in cultured spinal astrocytes. J Neurosci Res 91, 300–312. 10.1002/jnr.23144

[28] Pareek, T.K., Zipp, L., Letterio, J.J., 2013. Cdk5: An Emerging Kinase in Pain Signaling. Brain Disord Ther 2013, 003. 10.4172/2168-975X.S1-003

[29] Qu, Y.-J., Zhang, X., Fan, Z.-Z., Huai, J., Teng, Y.-B., Zhang, Y., Yue, S.-W., 2016. Effect of TRPV4-p38 MAPK Pathway on Neuropathic Pain in Rats with Chronic Compression of the Dorsal Root Ganglion. Biomed Res Int 2016, 6978923. 10.1155/2016/6978923

[30] R, S., W, R., Sp, H., Ah, D., 2004. Descending facilitatory control of mechanically evoked responses is enhanced in deep dorsal horn neurones following peripheral nerve injury. Brain research 1019. 10.1016/j.brainres.2004.05.108

[31] Roufayel, R., Murshid, N., 2019. CDK5: Key Regulator of Apoptosis and Cell Survival. Biomedicines 7, 88. 10.3390/biomedicines7040088

[32] S, F., 1999. Transmitters involved in antinociception in the spinal cord. Brain research bulletin 48. 10.1016/s0361-9230(98)00159-2

[33] Sun, J., Chen, F., Braun, C., Zhou, Y.-Q., Rittner, H., Tian, Y.-K., Cai, X.-Y., Ye, D.-W., 2018. Role of curcumin in the management of pathological pain. Phytomedicine 48, 129–140. 10.1016/j.phymed.2018.04.045

[34] Tawfik, V.L., Nutile-McMenemy, N., Lacroix-Fralish, M.L., Deleo, J.A., 2007. Efficacy of propentofylline, a glial modulating agent, on existing mechanical allodynia following peripheral nerve injury. Brain Behav Immun 21, 238–246. 10.1016/j.bbi.2006.07.001

[35] V, R., S, B., W, D., E, P., G, R., S, Vanvolsem, M, W., S, Vanneste, D, D.R., M, P., 2022. Real world data collection and cluster analysis in patients with sciatica due to lumbar disc herniation. Clinical neurology and neurosurgery 217. 10.1016/j.clineuro.2022.107246

[36] Wu, F., Miao, X., Chen, J., Liu, Z., Tao, Y., Yu, W., Sun, Y., 2013. Inhibition of GAP-43 by propentofylline in a rat model of neuropathic pain. Int J Clin Exp Pathol 6, 1516–1522.

[37] Wu, J., Wang, C., Ding, H., 2020. LncRNA MALAT1 promotes neuropathic pain progression through the miR-154-5p/AQP9 axis in CCI rat models. Mol Med Rep 21, 291–303. 10.3892/mmr.2019.10829

[38] Yan, P., Zhang, Z., Miao, Y., Xu, Y., Zhu, J., Wan, Q., 2019. Physiological serum total bilirubin concentrations were inversely associated with diabetic peripheral neuropathy in Chinese patients with type 2 diabetes: a cross-sectional study. Diabetol Metab Syndr 11, 100. 10.1186/s13098-019-0498-7

[39] Yang, L., Gu, X., Zhang, W., Zhang, J., Ma, Z., 2014. Cdk5 inhibitor roscovitine alleviates neuropathic pain in the dorsal root ganglia by downregulating N-methyl-D-aspartate receptor subunit 2A. Neurol Sci 35, 1365–1371. 10.1007/s10072-014-1713-9

[40] Yu, T., Zhang, X., Shi, H., Tian, J., Sun, L., Hu, X., Cui, W., Du, D., 2019. P2Y12 regulates microglia activation and excitatory synaptic transmission in spinal lamina II neurons during neuropathic pain in rodents. Cell Death Dis 10, 165. 10.1038/s41419-019-1425-4

[41] Yuan, Y., Zhen, L., Li, Z., Xu, Wenhua, Leng, H., Xu, Wen, Zheng, V., Luria, V., Pan, J., Tao, Y., Zhang, H., Cao, S., Xu, Y., 2020. trans-Resveratrol ameliorates anxiety-like behaviors and neuropathic pain in mouse model of post-traumatic stress disorder. J Psychopharmacol 34, 726–736. 10.1177/0269881120914221

[42] Zeb, A., Son, M., Yoon, S., Kim, J.H., Park, S.J., Lee, K.W., 2019. Computational Simulations Identified Two Candidate Inhibitors of Cdk5/p25 to Abrogate Tau-associated Neurological Disorders. Comput Struct Biotechnol J 17, 579–590. 10.1016/j.csbj.2019.04.010

[43] Zhang, H., Li, A., Liu, Y.-F., Sun, Z.-M., Jin, B.-X., Lin, J.-P., Yang, Y., Yao, Y.-X., 2023. Spinal TAOK2 contributes to neuropathic pain via cGAS-STING activation in rats. iScience 26, 107792. 10.1016/j.isci.2023.107792

[44] Zhang, Y.-H., Hu, H.-Y., Xiong, Y.-C., Peng, C., Hu, L., Kong, Y.-Z., Wang, Y.-L., Guo, J.-B., Bi, S., Li, T.-S., Ao, L.-J., Wang, C.-H., Bai, Y.-L., Fang, L., Ma, C., Liao, L.-R., Liu, H., Zhu, Y., Zhang, Z.-J., Liu, C.-L., Fang, G.-E., Wang, X.-Q., 2021. Exercise for Neuropathic Pain: A Systematic Review and Expert Consensus. Front Med (Lausanne) 8, 756940. 10.3389/fmed.2021.756940

[45] Zhang, Z., Leong, D.J., Xu, L., He, Z., Wang, A., Navati, M., Kim, S.J., Hirsh, D.M., Hardin, J.A., Cobelli, N.J., Friedman, J.M., Sun, H.B., 2016. Curcumin slows osteoarthritis progression and relieves osteoarthritis-associated pain symptoms in a post-traumatic osteoarthritis mouse model. Arthritis Res Ther 18, 128. 10.1186/s13075-016-1025-y

[46] Zhang, Z.-J., Zhao, L.-X., Cao, D.-L., Zhang, X., Gao, Y.-J., Xia, C., 2012. Curcumin inhibits LPS-induced CCL2 expression via JNK pathway in C6 rat astrocytoma cells. Cell Mol Neurobiol 32, 1003–1010. 10.1007/s10571-012-9816-4

[47] Zhao, W., Yan, J., Gao, L., Zhao, J., Zhao, C., Gao, C., Luo, X., Zhu, X., 2017. Cdk5 is required for the neuroprotective effect of transforming growth factor-β1 against cerebral ischemia-reperfusion. Biochem Biophys Res Commun 485, 775–781. 10.1016/j.bbrc.2017.02.130

[48] Zhao, Z., Li, X., Li, Q., 2017. Curcumin accelerates the repair of sciatic nerve injury in rats through reducing Schwann cells apoptosis and promoting myelinization. Biomed Pharmacother 92, 1103–1110. 10.1016/j.biopha.2017.05.099

